# Phosphoproteomic investigation of targets of protein phosphatases in EGFR signaling

**DOI:** 10.1101/2024.02.01.578427

**Authors:** Akihiro Eguchi, Jesper V. Olsen

## Abstract

Receptor tyrosine kinases (RTKs) initiate cellular signaling pathways, which are regulated through a delicate balance of phosphorylation and *de*phosphorylation events. While many studies of RTKs have focused on downstream-activated kinases catalyzing the site-specific phosphorylation, few studies have focused on the phosphatases carrying out the *de*phosphorylation. In this study, we analyzed six protein phosphatase networks using chemical inhibitors in context of epidermal growth factor receptor (EGFR) signaling by mass spectrometry-based phosphoproteomics. Specifically, we focused on protein phosphatase 2C (PP2C), involved in attenuating p38-dependent signaling pathways in various cellular responses, and confirmed its effect in regulating p38 activity in EGFR signaling. Furthermore, utilizing a p38 inhibitor, we classified phosphosites whose phosphorylation status depends on PP2C inhibition into p38-dependent and p38-independent sites. This study provides a large-scale dataset of phosphatase-regulation of EGF-responsive phosphorylation sites, which serves as a useful resource to deepen our understanding of EGFR signaling.

## Introduction

Site-specific protein phosphorylation is one of the most important post-translational modifications (PTMs) in cell signaling mediated by receptor tyrosine kinases (RTKs). It plays a central role in the phosphorylation cascades, a series of signaling events in which one kinase phosphorylates another, setting off a chain reaction that leads to the phosphorylation of thousands of proteins^1,2^. To date, much of the research in RTK signaling has focused on protein kinases that drive the cascade reaction. However, *de*phosphorylation events driven by protein phosphatases are also of considerable importance, as the signal amplification reaction in the phosphorylation cascade must be tightly regulated and eventually attenuated to avoid excessive phosphorylation^1,3^. Moreover, it is well-established that certain signaling proteins exhibit ligand-dependent *de*phosphorylation in response to ligand stimulation. This implies engagement of the molecular function of specific protein phosphatases that are not limited to the modulation of the phosphorylation cascade. Thus, comprehensive understanding of RTK signaling needs precise elucidation of the involvement and targets of protein phosphatases.

Since protein phosphatases generally target numerous phosphorylated substrates, large-scale analysis of protein phosphorylation states is needed to investigate the function of the phosphatases in cell signaling. Recent developments in mass spectrometry (MS)-based phosphoproteomics have made it possible to profile the cellular targets of protein phosphatases^4–10^. Kruse et al. applied phosphoproteomic strategies to identify substrates of protein phosphatase 2A on a global scale and outlined its *de*phosphorylation sequence motif preference^8^. Batth et al. and Vemulapalli et al. have recently examined the functions of the oncogenic protein tyrosine phosphatase SHP2 utilizing an allosteric inhibitor and MS-based phosphoproteomics to reveal biochemical activities of SHP2 in platelet-derived growth factor receptor and epidermal growth factor receptor (EGFR) signaling, respectively^9,10^.

Although numerous protein phosphatases are involved in EGFR signaling, phosphatase-centric phosphoproteomic analysis of EGFR signaling has been limited to focusing on eminent tyrosine phosphatases such as PTPN1^4^ and SHP2^10^. In this report, we utilized a protein phosphatase 2C (PP2C) inhibitor to investigate the phosphoproteome dynamics regulated by PP2C in EGFR signaling with comparison to five other phosphatases. For this, we profiled six phosphatase inhibitors: sanguinarine, Raphin1, KY-226, SHP099, NSC95397, and BCI, targeting PP2C, R15B-PP1c complex, PTPN1, SHP2, Cdc25, and DUSP1/6, respectively^11–16^. PP1c is also a serine/threonine phosphatase, PTPN1 and SHP2 are tyrosine phosphatases, and Cdc25 and DUSP1/6 are dual specific phosphatases. All phosphatases except R15B-PP1c have been reported to be involved in modulating the mitogen-activated protein kinase (MAPK) signaling pathway in different ways. SHP2 is the phosphatase whose functions in EGFR signaling have been most precisely investigated. SHP2 forms a complex with Gab1, Grb2, and SOS, and positively regulates the activation of MAPK pathway signaling; therefore, inhibiting SHP2 activity results in the down-regulation of EGF-dependent MAPK pathway signaling^17^. Conversely, DUSP family proteins are known as MAPK phosphatases (MKPs), and DUSP1 and DUSP6 *de*phosphorylate Erk to attenuate MAPK signaling in the nucleus and cytoplasm, respectively^18,19^. PTPN1 and Cdc25 have also been reported to target MAPKs or EGFR^20,21^.

PP2C is a member of the metal-dependent serine/threonine protein phosphatase family and has been reported to negatively regulate the MAPK signaling pathway in response to various cellular stimulation such as stress, interleukin, and transforming growth factor α stimulation, by *de*phosphorylating principal MAPK pathway kinase members such as TAK1, Erk, JNK, and p38^22–24^. However, its functions in EGFR signaling have scarcely been examined. Sanguinarine is a plant alkaloid that has been found to inhibit PP2C activity with specificity over other protein phosphatase families such as PP1, PP2A, and PP2B^11^. While sanguinarine has been used to investigate the activity of PP2C in various studies^25–27^, there has been no phosphoproteome-level examination of sanguinarine activity to date.

Our phosphoproteomics study examined the effects of each phosphatase inhibitor on EGF-dependent phosphorylation dynamics. We specifically focused on PP2C inhibition and summarized its effects on the p38-specific MAPK signaling pathway by classifying the PP2C inhibitor-regulated phosphosites into p38-dependent and p38-independent sites utilizing a p38-kinase inhibitor. Most of the identified p38-dependent sites are previously unassociated sites with p38 activity. This study offers a global and quantitative dataset of phosphatase-related EGF-dependent phosphorylation sites, which can serve as an information-rich resource to advance our understanding of EGFR signaling.

## Results

### Quantitative phosphoproteomics of EGFR signaling perturbed by six phosphatase inhibitors

To determine a favorable concentration of the inhibitors to be used for the ensuing proteomics experiment, we analyzed their concentration-dependent impact on cell toxicity and EGF-dependent phosphorylation of major protein kinases in EGFR signaling. We incubated Hela cells with 1 or 10 μM of each inhibitor for 15 min followed by stimulation with 20 nM EGF for 20 min, and evaluated the phosphorylation status of EGFR and downstream signaling proteins Akt and Erk by western blotting (Fig. S1). The treatment of 10 μM sanguinarine caused EGF-independent phosphorylation of all the detected sites. Moreover, all of the inhibitors showed cell toxicity at 10 μM (data not shown). On the other hand, we did not observe any change in baseline phosphorylation or cell toxicity by any inhibitor at 1 μM. Based on these results, we decided to use 1 μM or lower concentrations for further experiments. Of note, we reassuringly confirmed that phosphorylation of Erk was attenuated by inhibiting SHP2 due to its role as a positive regulator of the MAPK pathway signaling (Fig. S1).

Analyses of the site-specific phosphorylation states by MS-based quantitative phosphoproteomics was performed using HeLa cells treated with one of the inhibitors (0.1 or 1 μM) for 15 min followed by 8 or 20 min stimulation with EGF (20 nM) (Fig. 1A). Control samples included a no-stimulation control and an only-EGF (without any inhibitor) control. After cell lysis, 500 μg of proteins were applied to protein aggregation capture (PAC) with magnetic microparticles as carriers and subsequent on-bead digestion with an endoproteinase Lys-C/trypsin cocktail^28^, followed by phosphopeptide-enrichment by using magnetic Ti-IMAC beads. The resulting phosphopeptide mixtures were analyzed by liquid chromatography tandem mass spectrometry (LC-MS/MS) using the Evosep One LC coupled to an Orbitrap Exploris 480 mass spectrometer^29^. The mass spectrometer was operated in data-independent acquisition (DIA)-mode utilizing 60 samples-per-day (SPD) LC gradient.

**Figure 1.**
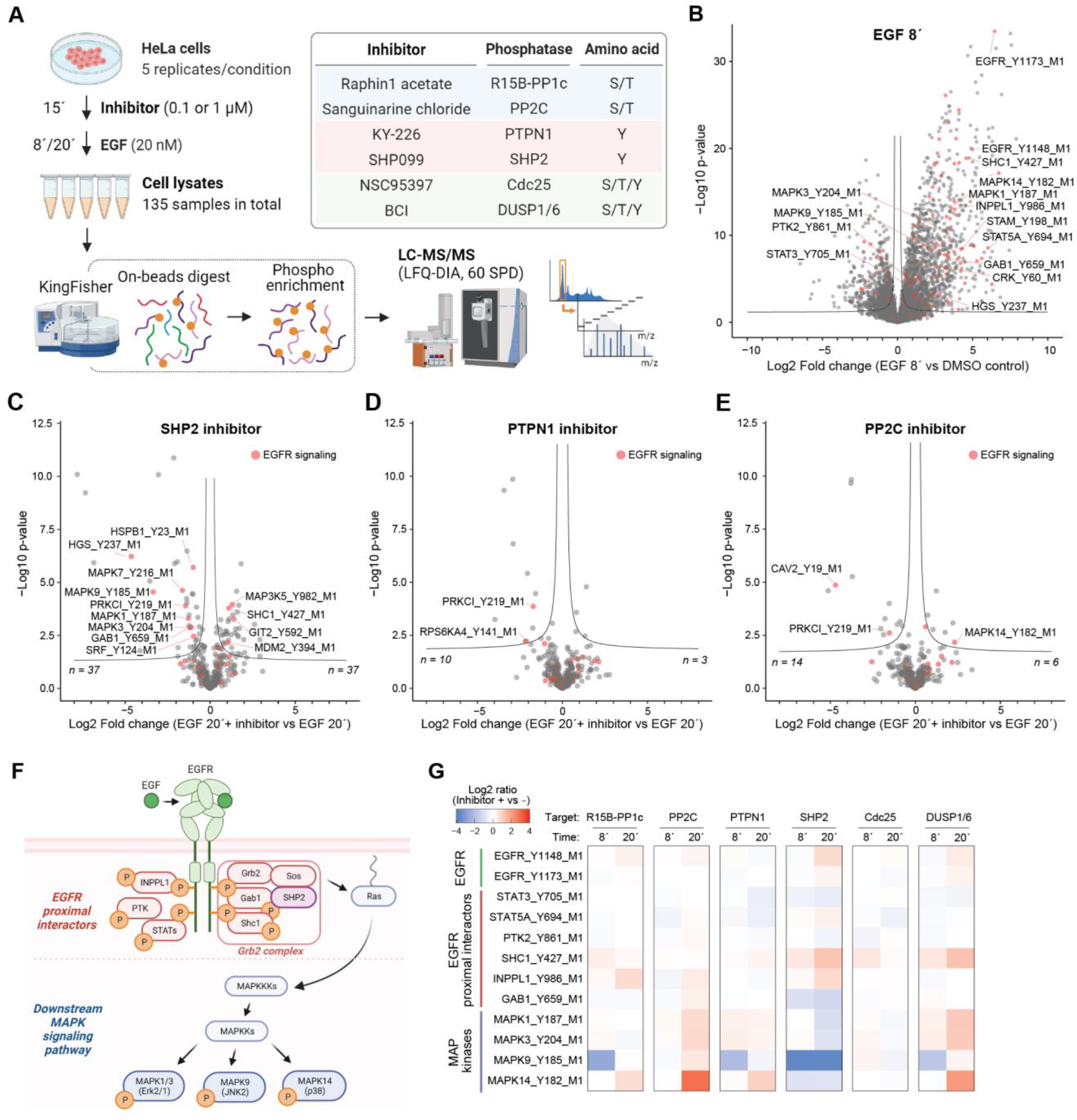
MS-based phosphoproteomics of EGF signaling with treatment of six inhibitors targeting protein phosphatases. (A) Schematic illustration of MS-based phosphoproteomics sample preparation and a list of protein phosphatase inhibitors used. The figure was created using BioRender.com. (B-E) Volcano plots highlighting EGFR signaling-related proteins (red). (B) Fold change represents EGF (20 nM, 8’
s) treatment versus no-stimulation control. (C–E) Fold change represents EGF (20 nM, 20’) with versus without the inhibitor of either (C) SHP2, (D) PTPN1, or (E) PP2C. (F) Illustration of EGF-dependent change of protein interaction and phosphorylation. The figure was created using BioRender.com. (G) Heatmap showing the log2-transformed peptide abundance changes induced by inhibitor treatment.

Each experimental condition was analyzed with five biological replicates, and we collectively identified 31,738 classI phosphosites^30^ in total (Fig. S2A: 5.8% pY: 1,831, 23.1% pT: 7,331, 71.1% pS: 22,576). For label-free quantitation (LFQ), we initially filtered the phosphosites based on the number of missing values and required at least 70% valid quantitation values in each experimental condition. The matrix of LFQ-based intensities for the resulting 13,718 phosphosites were normalized and imputed. The peptides were further filtered based on the number of missing values in only-EGF condition, and resulting 7,795 phosphosites were used for the further analysis. Comparing the EGF-only and no-stimulation conditions using a volcano plot displaying the log2 fold-change (FC) differences on x-axis and negative log10-transformed t-test significance value (p-value) on y-axis, manifested up-regulation of lots of phosphosites (Fig. 1B). These includes regulatory tyrosine phosphorylation sites on EGFR itself and EGFR signaling-related proteins such as SHC1 (Y427), INPPL1 (Y986), STAT5A (Y694), GAB1 (Y659), MAPK1 (Y187), MAPK3 (Y204), and MAPK14 (Y182), confirming that our data successfully capture known phosphoproteome changes triggered by EGFR activation.

To dissect the global effects of the phosphatase inhibition, we initially focused on the regulation of tyrosine phosphorylation sites. The volcano plot comparing SHP2 inhibitor (+) and (−) condition showed many regulated sites including down-regulation of MAPK1 (Y187), MAPK3 (Y204), and GAB1 (Y659), confirming previous reports^10^ (Fig. 1C). Conversely, the inhibition of another tyrosine phosphatase PTPN1 did not result in inducing a noticeable change of global tyrosine phosphorylation status (Fig. 1D), substantiating the distinct importance of SHP2 in EGFR signaling. Similarly, inhibition of PP2C also did not result in substantial change in global tyrosine phosphorylation status. However, the up-regulation of the Y182 site of MAPK14/p38α, located in its activation loop, suggested that PP2C is involved in the regulation of the p38α activity in EGFR signaling (Fig. 1E). The identified EGFR signaling-related proteins that undergo tyrosine phosphorylation can be categorized into EGFR proximal interactors and downstream MAPK signaling pathway proteins (Fig. 1F). EGFR proximal interactors are proteins that are either directly recruited to the activated receptor or proteins that form a complex with other proteins on EGFR. The protein complex represented by Grb2, which includes SHP2, plays a pivotal role in triggering the MAPK pathway signaling by activating Ras. We assessed the contribution of each phosphatase inhibitor on the phosphorylation of those proteins by plotting log2 FC of phosphosite abundance changes induced by the inhibitor treatment (Fig. 1G). Two phosphorylation sites of EGFR (Y1148 and Y1173) showed almost no or slight increase of phosphorylation at 20 min by most of the inhibitors. Among EGFR proximal interactors, phosphorylation of SHC1 (Y427) and INPPL1 (Y986) were generally more susceptible to changes than other sites were. Activating phosphorylation in the kinase activation loop of four identified MAPKs exhibited similar trends: down-regulation by the SHP2 inhibition and up-regulation by inhibition of PP2C, PTPN1, and DUSP1/6. Notably, phosphorylation of MAPK14 (Y182) was up-regulated by four inhibitors, and the PP2C inhibitor induced the largest up-regulation (Fig. 1G).

Next, we focused on phosphatase-regulation of phosphorylation of serine and threonine sites. We analyzed phosphosites showing up-regulation by EGF treatment at either time point and compared their phosphorylation changes induced by the SHP2 inhibitor and the PP2C inhibitor (Fig. S2B and S2C). The global trends depicted by the volcano plots showed regulation in the opposite direction: down-regulation by the SHP2 inhibitor and up-regulation by the PP2C inhibitor. Of note, we found the up-regulation of S439 site on MAP3K7/TAK1, an upstream MAPK kinase kinase of p38α and a known direct target of PP2C, despite that the regulation was not significant. We summarized the effect of the phosphatase inhibition on serine/threonine phosphorylation sites on EGFR signaling-related proteins in the heatmap (Fig. S2D). The phosphorylation sites in clusters A and B displayed the same regulation pattern as MAPK14/p38α (Y182) shown in Fig. 1G, presenting up-regulation by the PP2C inhibitor and down-regulation by the SHP2 inhibitor. Therefore, these peptides are potential candidates for substrates in the p38 kinase signaling pathway, and indeed, the clusters included phosphorylation sites of HSPB1 (S15, S78, S82), which are known substrate of MAPKAPK2, a direct downstream kinase substrate of the p38 kinase. From these findings, we deemed that our data adequately capture the rewiring of intracellular signaling networks caused by inhibition of the phosphatases.

### Cell cycle- and mitosis-related proteins are major targets of phosphatases upon dephosphorylation events in EGFR signaling

To explore the impact of phosphatase regulation in EGF-dependent signaling in more detail, we next focused on the phosphosites showing EGF-dependent down-regulation to investigate the role of phosphatases on *de*phosphorylation events triggered by EGFR activation. Dephosphorylated peptides were extracted based on fold change (FC < 1/2) and significance (p-value < 0.05) comparing the EGF-only and no-stimulation conditions for each time point (Fig. 2A; 441 peptides for 8 min, 392 for 20 min conditions, and 210 peptides shared). To functionally interpret the global tendency of EGF-dependent *de*phosphorylated sites, we performed an iceLogo^31^-based sequence motif enrichment analysis to characterize the preferences of amino acids surrounding the phosphorylation sites (±7) (Fig. S3). This analysis revealed an overrepresentation of negatively charged amino acids such as aspartic acid and glutamic acid at most positions close to the down-regulated phosphorylation sites, especially +1 and +3 positions, which is a known signature of casein kinases^32^. This result implies the *de*phosphorylation events are driven by down-regulation of casein kinases or up-regulation of certain phosphatases that prefer negatively charged amino acids surrounding their target phosphorylation sites. Principal component analysis (PCA) of the shared 210 phosphosites clearly separated the treatments by PC1 and treatment time by PC2 with the SHP2 inhibitor (+) conditions showing the largest deviation from the control conditions at both time points (Fig. 2B).

**Figure 2.**
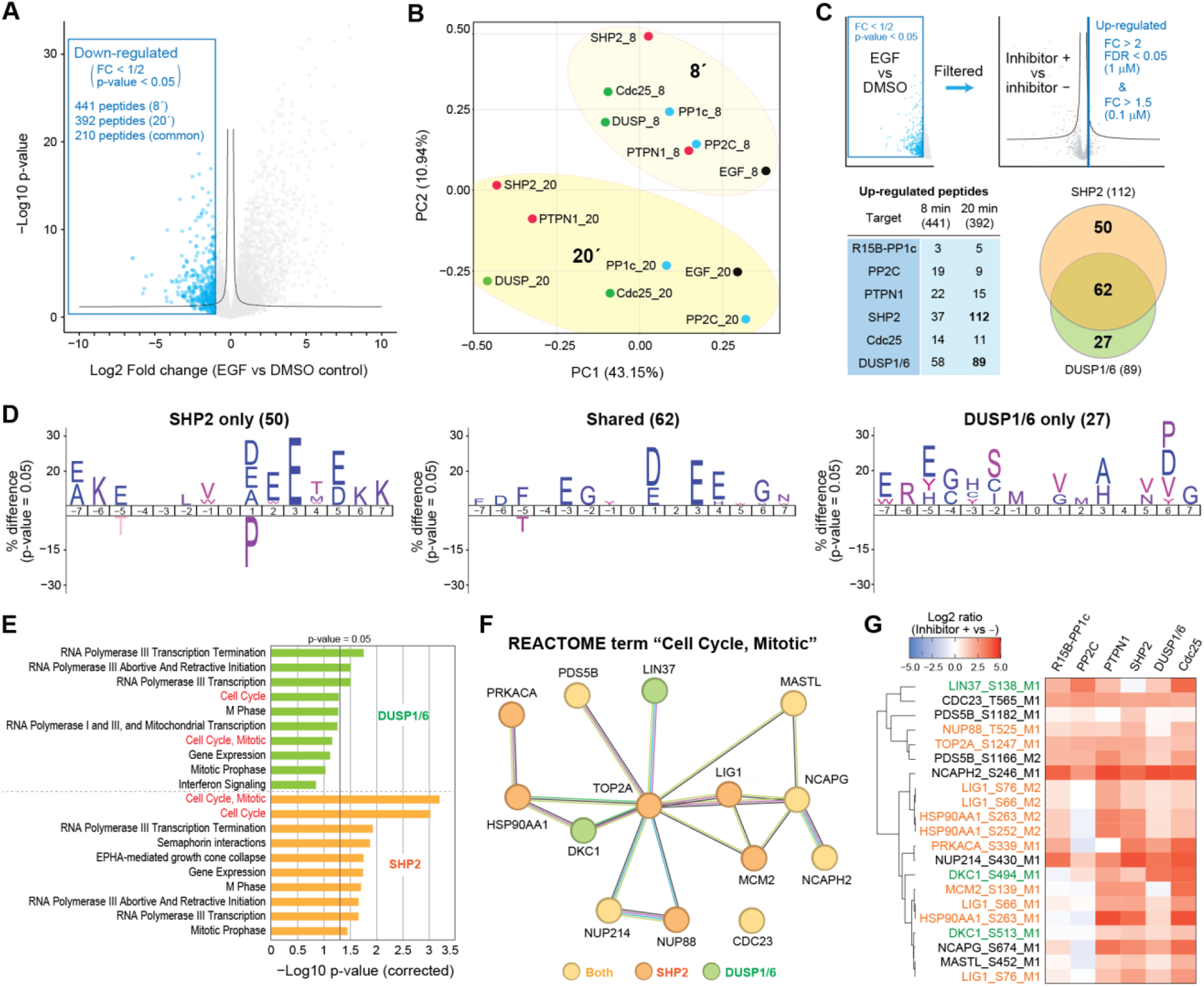
Investigation of phosphosites showing EGF-dependent dephosphorylation. (A) Volcano plot highlighting phosphosites showing EGFR-dependent down-regulation. (B) PCA of the log2-transformed abundance of phosphosites down-regulated by both 8 and 20 min of EGF incubation. (C) Scheme to filter peptides showing up-regulation by inhibitor treatment and summary of the numbers of the up-regulated phosphosites for each inhibitor (D) Sequence motif enrichment analysis of 15 residues surrounding the regulated phosphosites. (E) Reactome enrichment analysis using InnateDB using up-regulated phosphosites. (F) STRING-based protein network of the identified proteins involved with cell cycle and mitosis. (G) Heatmap showing the log2-transformed phosphosite abundance changes induced by inhibitor treatment.

To investigate the propensity of the phosphorylation site regulation by each phosphatase, we compared the phosphosites that showed attenuated down-regulation by treatment of each inhibitor (Fig. 2C; FC > 2 and FDR < 0.05 for 1 μM, FC > 1.5 for 0.1 μM). Among the six inhibitors, the SHP2 and DUSP1/6 inhibitors gave higher number of regulated peptides than the other four inhibitors especially at 20 min time point, and a large part of the regulated peptides were common between the two conditions. Sequence motif enrichment analysis was performed using these phosphosites to identify possible sequence preferences of each phosphatase with respect to the amino acid sequence surrounding its target sites (Fig. 2D). Phosphosites regulated only by the SHP2 inhibitor or by both showed enrichment of the consensus motif of casein kinases, while sites regulated only by the DUSP1/6 inhibitor showed no obvious consensus motifs. Furthermore, inhibition of SHP2 negatively enriches proline in the +1 position, which is important for MAPKs substrate recognition, consistent with SHP2 inhibition leading to downregulation of MAPKs.

Reactome enrichment analysis by InnateDB^33^ using a list of phosphoproteins regulated by the SHP2 or DUSP1/6 inhibitor treatment showed that RNA polymerase-related proteins and cell cycle/mitosis-related proteins were most significantly affected in both conditions (Fig. 2E). In particular, Reactome terms “Cell Cycle” and “Cell Cycle, Mitotic” showed substantial difference between the two inhibitors. In our dataset, a total of 14 phosphoproteins associated with these terms were identified, and 6 out of 14 were common between the two inhibitors (Fig. 2F). A heatmap of these cell cycle protein phosphorylation sites showed that most of these sites are upregulated by inhibition of most phosphatases (Fig. 2G). This suggests that various phosphatases are involved with regulation of phosphorylation status of mitosis-related proteins in response to EGFR activation.

### PP2C in particular shows regulatory function in p38 signaling cascade

Next, we investigated the involvement of the phosphatases in regulation of phosphorylation cascades in EGFR signaling, focusing on phosphosites exhibiting EGF-dependent up-regulation (FC > 1.5) for each time point (Fig. 3A; 1,171 peptides at 8 min, 1,153 at 20 min, and 890 peptides are shared). PCA of the shared 890 phosphosites clearly differentiated the two time points by PC1 and the inhibitors by PC2. The distribution by inhibitor treatment was more scattered at 20 min than at 8 min, and the SHP2 inhibitor showed the largest deviation from the control condition (Fig. 3B). This is consistent with the fact that SHP2 plays an important and unique role for regulating EGF-dependent activation of the MAPK pathway signaling. Interestingly, only the PP2C inhibitor showed a shift in the opposite direction from the other five inhibitors.

**Figure 3.**
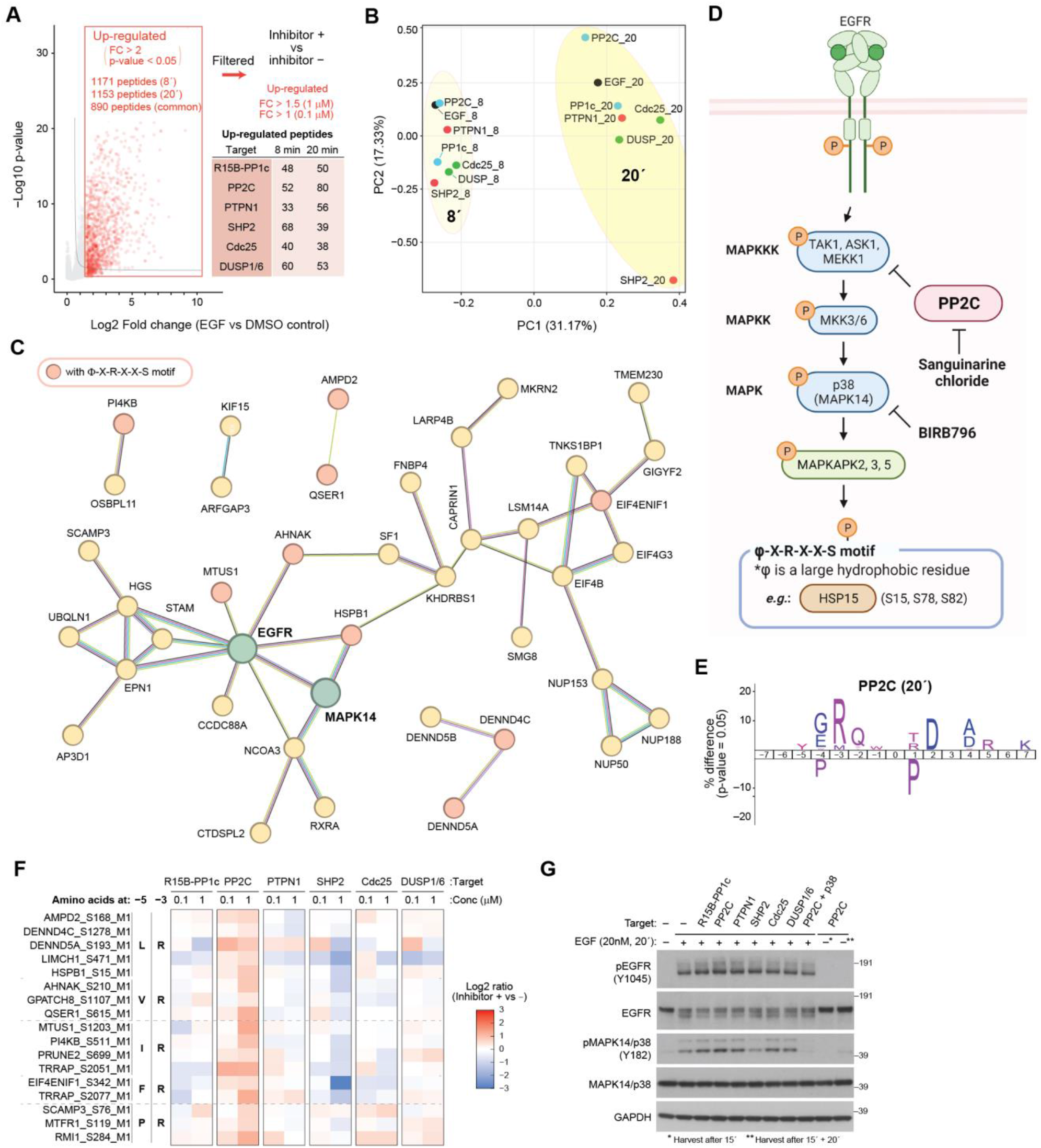
Investigation of phosphosites showing EGF-dependent up-regulation. (A) Scheme to filter peptides showing up-regulation by inhibitor treatment and summary of the numbers of the up-regulated phosphosites for each inhibitor (B) PCA of the log2-transformed abundance of phosphosites up-regulated by both 8 and 20 min of EGF incubation. (C) STRING-based protein network of proteins showing sanguinarine-dependent up-regulation of its phosphorylation. Proteins with at least one connection with other proteins are included. (D) Illustration of the p38 pathway activated by EGFR. The figure was created using BioRender.com. (E) Sequence motif enrichment analysis of 15 residues surrounding the regulated phosphosites. (F) Heatmap showing the log2-transformed phosphosite abundance changes induced by inhibitor treatment. See also Fig. S4 for the heatmap containing all 80 peptides. (G) Western blotting analysis using the cell lysates of Hela cells pre-treated with the inhibitors (500 nM) for 15 min followed by 20 min incubation with EGF (20 nM).

Similar to the analysis of down-regulated phosphosites depicted in Fig. 2A, we summarized the numbers of the phosphosites that showed enhanced up-regulation by the treatment of each inhibitor (Fig. 3A; FC > 1.5 for 1 μM and FC > 1 for 0.1 μM). We focused on 80 phosphosites which was regulated by PP2C inhibition at 20 min and built a functional phosphoprotein network based on STRING^34^ with EGFR included (Fig. 3C). Proteins highlighted in red color have a regulated phosphorylation site with a Φ-X-R-X-X-S motif, where Φ is a large hydrophobic amino acid residue. This is the established recognition motif of MAPKAPK2^35,36^, a direct downstream kinase of the p38 kinase (Fig. 3D). Indeed, sequence motif enrichment analysis of the 80 regulated phosphorylation sites exhibited an enrichment of arginine at –3 position (Fig. 3E), and 17 sites out of 80 had an arginine at –3 position and a hydrophobic amino acid (L, V, I, F, or P) at –5 position. The heatmaps plotting log2 FC of phosphosite abundance changes induced by inhibitor treatment showed that most of them were down-regulation by SHP2 inhibition (Fig. 3F, also see Fig. S4 for all the 80 peptides), implying that these phosphosites are indeed downstream target of MAPK pathway signaling. These findings were also confirmed by western blotting using the cells stimulated with EGF in the presence of the inhibitors (Fig. 3G, lanes 1 to 8). Only the SHP2 inhibitor inhibited phosphorylation of p38α (Y182), whereas the other five inhibitors caused increase in the phosphorylation to some extent. Among the five inhibitors, the PP2C inhibitor showed the strongest increase, and the phosphorylation was almost completely blocked by addition of the p38 inhibitor BIRB796 (Fig. 3G, lane 9). In addition, we confirmed that the PP2C inhibitor induced minimum background phosphorylation of p38α in absence of EGF after 15 min incubation or even after additional 20 min incubation (Fig. 3G, lanes 10 and 11).

To ascertain whether these proteins are genuinely regulated in a p38α-dependent or -independent manner, we performed another phosphoproteomics experiment using cells stimulated with EGF in the presence of sanguinarine and BIRB796 (Fig. 4A). In this experiment, we used different concentrations (5, 50, and 500 nM) of sanguinarine to obtain information of its concentration-dependent response. After data filtration, we identified 9,831 class I phosphosites, which included 54 out of 80 phosphosites regulated by sanguinarine in Fig. 3. The functional protein network depicted in Fig. 4B keeps the proteins identified in the second-round analysis, and 7 out of 9 proteins with a Φ-X-R-X-X-S motif were retained. Unsupervised hierarchical clustering of these phosphosites based on their regulation at 20 min divided the sites into three clusters (Fig. 4C). The sites in cluster 1 showed down-regulation in the presence of BIRB796, and 5 out of 7 proteins with a Φ-X-R-X-X-S motif were in this cluster. This BIRB796-responsive group included not only HSPB1 (S15), a known target of MAPKAPK2, but also several previously uncharacterized p38-dependent phosphosites such as LSM14A (S183), AMPD2 (S168), AHNAK (S210), and QSER1 (S615). These phosphoproteins are potential candidates for p38 downstream target proteins. The BIRB796-unaffected peptides in the cluster 3 included PI4KB (S511) and OSBPL11 (S189), previously discovered as EGF-dependent phosphorylation sites^37^. We identified them as PP2C-dependent but p38-independent phosphorylation sites. They were observed in the protein network in Fig. 3C and 4B, but they do not have connection with any p38-related proteins. These findings suggests that these sites are phosphorylated by other kinases, or possibly directly *de*phosphorylated by PP2C, despite that S511 site of PI4KB has a recognition motif of MAPKAPK2.

**Figure 4.**
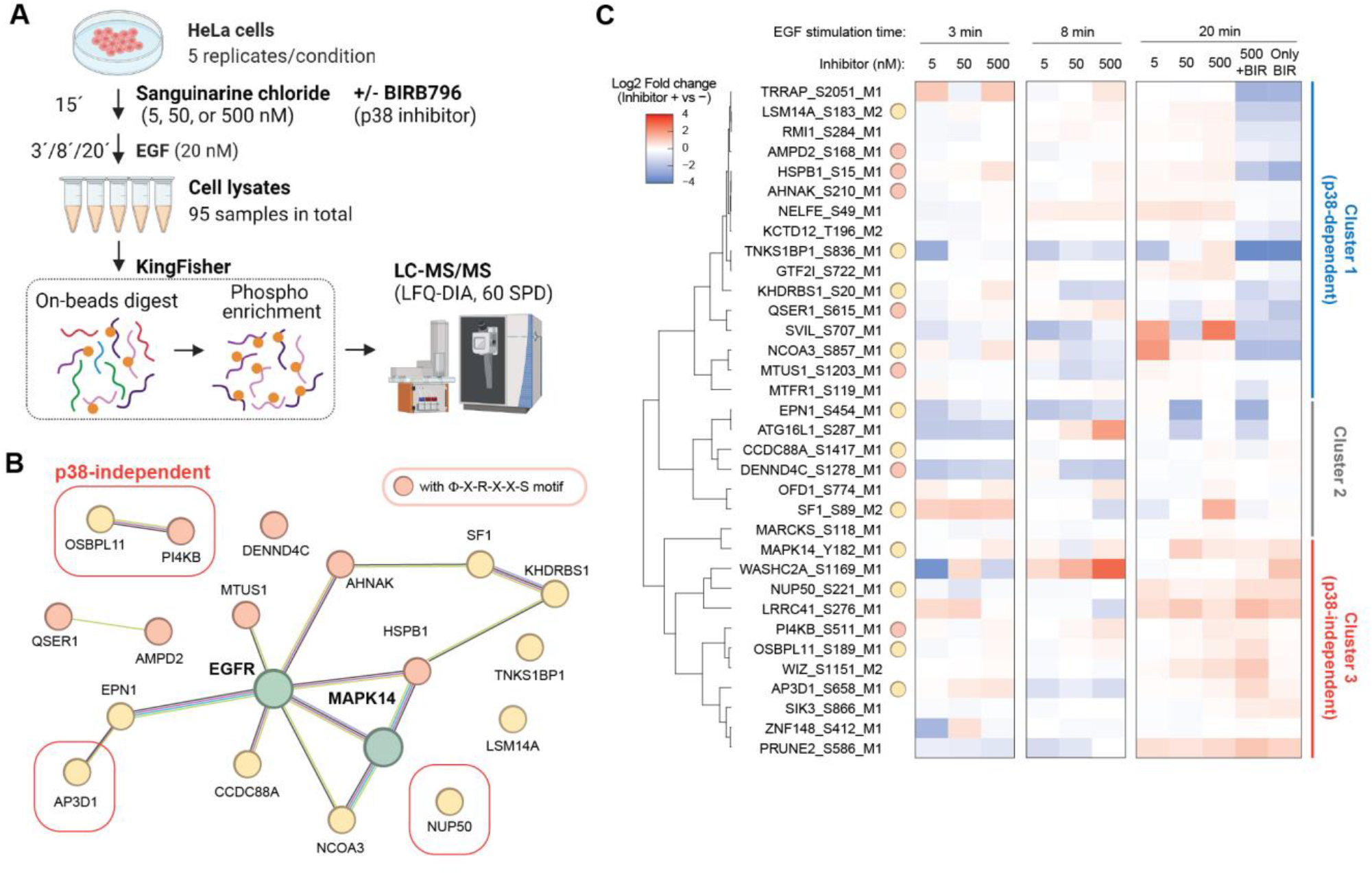
MS-based phosphoproteomics of EGF signaling with treatment of PP2C inhibitor sanguinarine in presence of p38 inhibitor BIRB796. (A) Schematic illustration of MS-based phosphoproteomics sample preparation. The figure was created using BioRender.com. (B) STRING-based protein network of proteins showing sanguinarine-dependent up-regulation of its phosphorylation. (C) Heatmap showing the log2-transformed phosphosite abundance changes induced by inhibitor treatment.

## Discussion

In this study, we used six different phosphatase inhibitors to comprehensively examine protein phosphorylation states affected by protein phosphatases in context of EGFR signaling. An important aspect of research targeting protein phosphatases is the selection of the optimal experimental system, *e*.*g*., whether to perform *in vitro*^38^ or *in cellulo*^5–10^, and whether to use inhibitors^7–10^ or silencing of targets^4–6^ such as siRNA knockdown for phosphatase inhibition. To investigate the role of protein phosphatase in dynamic cellular functions such as cell signaling, it is necessary to use an *in cellulo* system and limit the time during which phosphatase function is restricted to a short period. Consequently, in this study, we performed MS-based phosphoproteomics using EGF-stimulated cells in the presence of small chemical inhibitors to investigate the role of protein phosphatases in EGFR signaling. On the other hand, it should be noted that it is difficult to distinguish between direct and indirect targets of phosphatases when analyzing *in cellulo* systems. This is because phosphatase-mediated *de*phosphorylation affects the activity of other kinases and phosphatases. It needs to be confirmed whether the phosphorylation sites found in this study are direct substrates of the phosphatases, *e*.*g*., by *in vitro* assay using phosphorylated peptides. Moreover, this study used small molecule inhibitors, the specificity of which must be carefully considered. Sanguinarine, for example, is known to inhibit AMPK and Na+/K+-ATPase at higher concentration^39,40^. Although these effects were minimized in this study using concentrations below 1 μM, interpreting the data requires some caution.

The effects of PP2C on p38 signaling have been reported in various cellular activities other than EGFR signaling, and we confirmed that a similar function is also found in EGFR signaling and that its contribution is greater than that of many other MAPK-related phosphatases. Our analysis included several phosphatases that have been known to be associated with regulation of the activity of MAPKs, and their inhibition certainly resulted in up-regulation of p38 phosphorylation. However, only PP2C inhibition affected phosphorylation of proteins downstream of p38. This suggests that PP2C plays a particularly important role in p38 signaling, possibly because PP2C also targets an upstream kinase TAK1, contributing to stronger p38 signaling inhibition as observed.

As for their involvement in EGF-dependent *de*phosphorylation, most of the phosphatases targeted in this study had effects on mitosis-related proteins. Phosphorylation of mitosis-associated proteins is known to be regulated by very large stoichiometric ratios, and the abundance of phosphorylated sites present in the deactivated state is very low^41^. Our findings suggest the possibility that various phosphatases are involved in the *de*phosphorylation of these proteins, keeping phosphorylated populations at very low stoichiometry.

As a potent inhibitor, sanguinarine has been used in various studies to investigate the biological role of PP2C. This work provides the first global phosphoproteome-level investigation of the effects of sanguinarine treatment. This also applies to the other three inhibitors with the exception of SHP099^9,10^ and BCI^42^, for which phosphoproteomic studies have already been reported. This is also the first phosphoproteomics study on BCI in terms of its relationship to EGFR signaling. The wide concentration range (5, 50, 100, 500, and 1000 nM) and EGF stimulation time (3, 8, and 20 min) of sanguinarine, in particular, are expected to provide useful information for further inhibitor studies.

In summary, this study provides a huge resource on perturbation of EGF-dependent phosphorylation sites under conditions of phosphatase inhibition utilizing small molecule inhibitors and demonstrates the utility of this approach for future phosphatase-centric EGFR signaling analysis.

## Method

### Materials

Raphin1 acetate (HY-123960A), Sanguinarine chloride (HY-N0052A), KY-226 (HY-120327), SHP099 (HY-100388), NSC95397 (HY-108543), and BCI (HY-115502) were purchased from MedChemExpress. BIRB796 (Axon 1358) was purchased from Axon Medchem. Recombinant human EGF (AF-100-15) was purchased from Peprotech.

### Cell Culture

HeLa cells were cultured at 37°C in a humid atmosphere with 5% CO2 in DMEM supplemented with 10% heat-inactivated fetal bovine serum (FBS) and penicillin/streptomycin. The medium without FBS was used as starvation medium.

### Cell lysis and western blotting

HeLa cells (2.0 × 10^5^ cells/well) were seeded in 6-well plates, incubated overnight, and the medium was replaced by starvation medium. After the overnight starvation, cells were treated with indicated concentration of an inhibitor for indicated time, followed by stimulation with 20 nM EGF for indicated time. After the stimulation, cells were washed with PBS, lysed using modified RIPA buffer (50 mM Tris-HCl (pH7.5), 150 mM NaCl, 1% NP-40, 0.1% sodium deoxycholate, 1 mM ethylenediaminetetraacetic acid, 5 mM β-glycerophosphate, 5 mM NaF, 1 mM sodium orthovanadate) supplemented with a tablet of protease inhibitor cocktail (Merck), scraped, centrifuged at 12,000 × g for 20 min at 4°C, and the supernatants were used for the further procedure. Protein concentration was determined using a BCA assay kit (Thermo Fisher Scientific), and the volume was adjusted so that the same amount of proteins would be applied to SDS Polyacrylamide gel electrophoresis (SDS-PAGE). Cell lysates were mixed with NuPAGE LDS Sample Buffer (4×) (Thermo Fisher Scientific) supplemented with 100 mM dithiothreitol, boiled for 10 min at 70°C, separated by SDS-PAGE, and transferred to PVDF membranes. Membranes were blocked with 5% BSA/PBS-T, incubated with indicated primary antibody overnight at 4°C, and incubated with HRP-conjugated secondary antibody for 1h at ambient temperature, and developed using Novex™ ECL Chemiluminescent Substrate Reagent Kit (Thermo Fisher Scientific). Primary antibodies used were rabbit anti-EGFR (ab32198; Abcam), rabbit anti-phospho-EGFR (pY1045) (#2237; Cell Signaling Technology), mouse anti-phospho-EGFR (pY1068) (#2236; Cell Signaling Technology), rabbit anti-Erk1/2 (#4695; Cell Signaling Technology), mouse anti-phospho-Erk1/2 (T202/Y204) (#9106; Cell Signaling Technology), rabbit anti-Akt (#9272; Cell Signaling Technology) rabbit anti-phospho-Akt (S473) (#9271; Cell Signaling Technology), rabbit anti-p38 MAPK (#9212; Cell Signaling Technology), rabbit anti-phospho p38 MAPK (T180/Y182) (#9211; Cell Signaling Technology) and mouse anti-GAPDH (Abcam).

### Sample preparation for phosphoproteomics

HeLa cells were seeded in 15 cm dishes, incubated overnight, and the medium was replaced by starvation medium. After the overnight starvation, cells were treated with indicated concentration of an inhibitor for indicated time, followed by stimulation with 20 nM EGF for indicated time. After the stimulation, cells were washed with PBS, lysed using modified RIPA buffer supplemented with a tablet of protease inhibitor cocktail, scraped, centrifuged at 12,000 × g for 20 min at 4°C, and the supernatants were used for the further procedure. Protein concentration was determined using a BCA assay kit (Thermo Fisher Scientific), and 500 μg of proteins was reduced and alkylated with 5 mM TCEP and 10 mM CAA for 15 min at ambient temperature. Peptides were digested using PAC method and eluted in 50 mM TEAB (pH 8.0) containing 0.5 μg of Lys-C and 1 μg of Trypsin. After the digestion overnight at 37°C, supernatants were acidified with 10% FA, loaded to Sep-Pak tC18 96-well μElution Plate for desalting, and eluted with 40% ACN and 60% ACN subsequently. Eluants were adjusted to 80% ACN/5% TFA/1 M glycolic acid (GA) and incubated with 20 μL of Ti-IMAC beads. The beads were subsequently washed with 80% ACN/5% TFA/1 M GA, 80% ACN/1% TFA, and 10% ACN/0.2% TFA, and enriched phosphosites were eluted in 1% NH3. The eluents were acidified with 10% TFA, filtered, and loaded onto Evotips for LC-MS/MS analysis.

### LC-MS/MS and MS data analysis

Samples for all the phosphoproteomics analysis were analyzed using the Evosep One system coupled with Orbitrap Exploris 480 MS (Thermo Fisher Scientific) in data-independent acquisition (DIA) mode. Peptides were eluted in either an in-house packed 15 cm, 150 mm i.d. capillary column with 1.9 mm Reprosil-Pur C18 beads (Dr. Maisch) with a heater set to 60°C a performance column (EV1137, EvoSep) with a heater set to 40°C, and 60 samples per day were analyzed using a preprogrammed gradient. The spray voltage was set to 2 kV, funnel RF level to 40, and capillary temperature to 275°C. Full MS resolution was set to 120,000 at m/z 200, full MS AGC target to 300%, and IT to 45 ms. Mass range was set to 350–1400. AGC target for fragment spectra set to 100%. 16 windows of 39.5 m/z scans from 472 to 1143 m/z with 1 m/z overlap were used. Resolution was set to 45,000 and IT to 86 ms. Normalized collision energy was set to 27%. All data were acquired in profile mode using positive polarity.

All MS raw files were analyzed by Spectronaut (Biognosys) v18.5 in a library-free mode (direct DIA+) using the human database (20,422 entries from UP000005640_9606 with signal peptides removed, reviewed 2023) supplemented with a database of common contaminants. Carbamidomethylation of cysteine was considered a fixed modification, and acetylation of the protein N-terminus, oxidation of methionine, and phosphorylation of serine, threonine, and tyrosine were considered variable modifications. Maximum number of variable modifications were set at 3, and minimum localization threshold was set at 0.75. False discovery rate (FDR) for Peptide spectrum match (PSM), peptide, and protein groups was set at 0.01. Cross-run normalization was turned off.

### Bioinformatics

Phosphosites quantified at least 70% of the replicates in at least one condition were used for the further analysis. The abundance values were log2-transformed, and normalized using the “normalizecyclicloess” function of the “limma” package in R. The missing values in the dataset were categorized into “partially observed values (POV)” and “missing in the entire condition (MEC)” and imputed using the “wrapper.impute.slsa” function and the “wrapper.impute.detQuant” function of “DAPAR” package in R, respectively. Batch correction was performed using Combat based on the experimental days. Volcano plots were depicted using Perseus software with a setting of FDR = 0.05 and s0 value = 0.1. “EGFR signaling” proteins consist of proteins that has KEGG terms of “ErbB signaling pathway”, “Endocytosis”, “MAPK signaling pathway”, “Jak-STAT signaling pathway”, or “Phosphatidylinositol signaling system”.

## Acknowledgement

Work at The Novo Nordisk Foundation Center for Protein Research (CPR) is funded in part by a donation from the Novo Nordisk Foundation (NNF14CC0001). This project was funded by the European Union’s Horizon 2020 research and innovation program under grant agreement EPIC-XS-823839 and ERC synergy grant 810057-HighResCells.

## Author contributions

A.E. performed all the experiments and MS data analysis, supervised by J.V.O. A.E. and J.V.O. wrote and approved the manuscript.

## Figures

**Figure S1.**
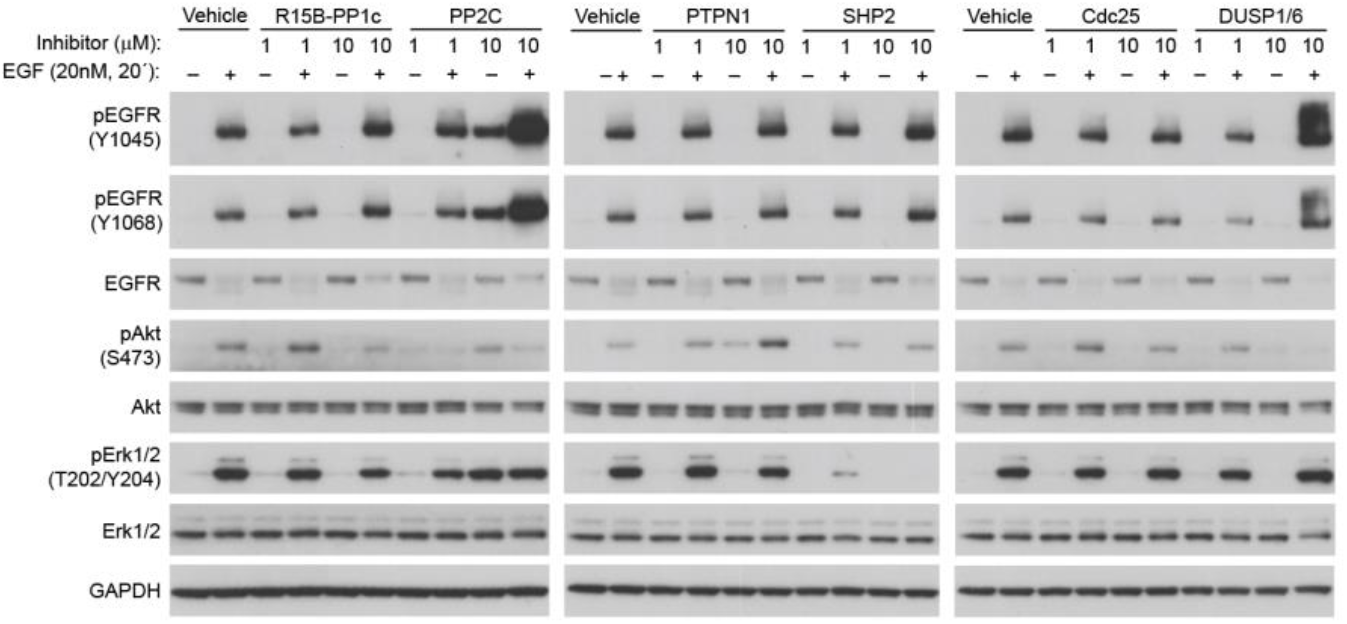
Western blotting analysis using the cell lysates of Hela cells pre-treated with the inhibitors (0.1 or 1 μM) for 15 min followed by 20 min incubation with EGF (20 nM).

**Figure S2.**
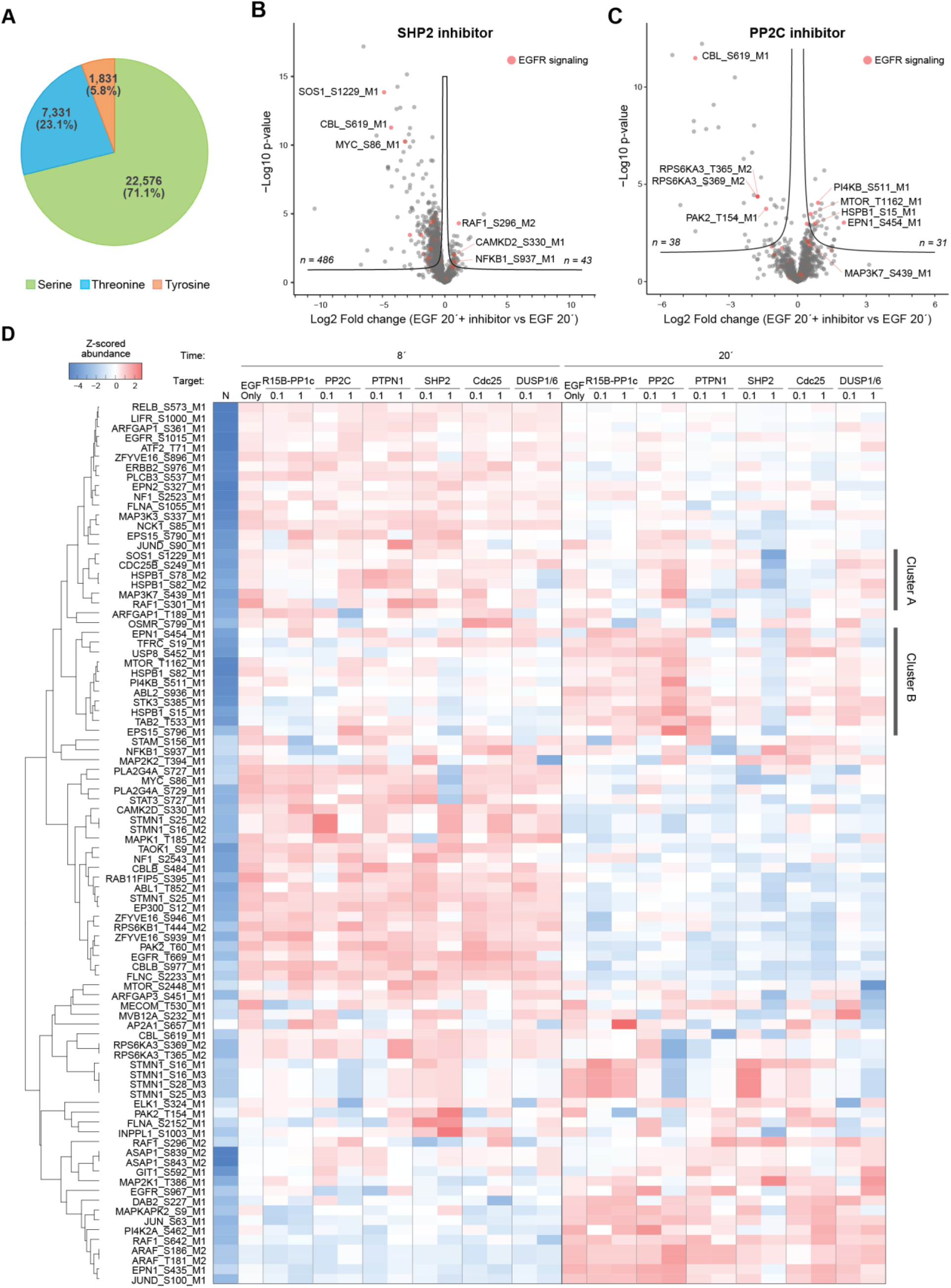
(A) The numbers and distribution of phosphorylated amino acids. (B, C) Volcano plot highlighting EGFR signaling-related proteins (red). Fold change represents EGF (20 nM, 20’
s) with the inhibitor of either (B) SHP2 or (C) PP2C. (D) Heatmap showing the z-scored phosphosite abundance.

**Figure S3.**
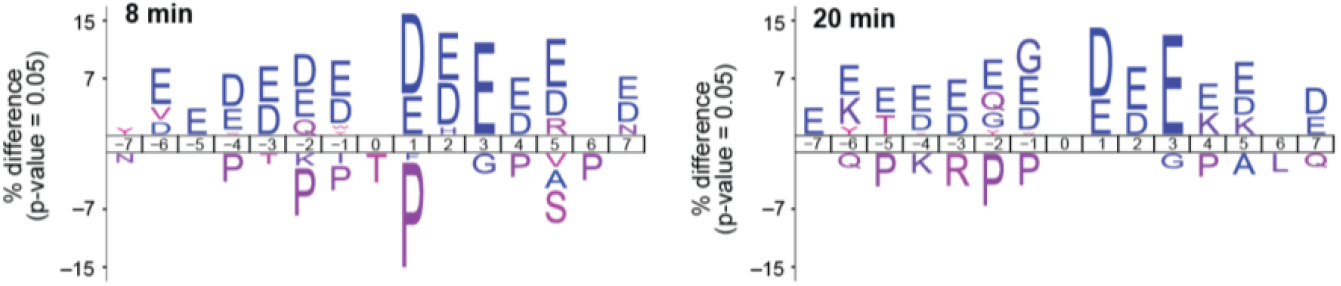
Sequence motif enrichment analysis of 15 residues surrounding the regulated phosphosites.

**Figure S4.**
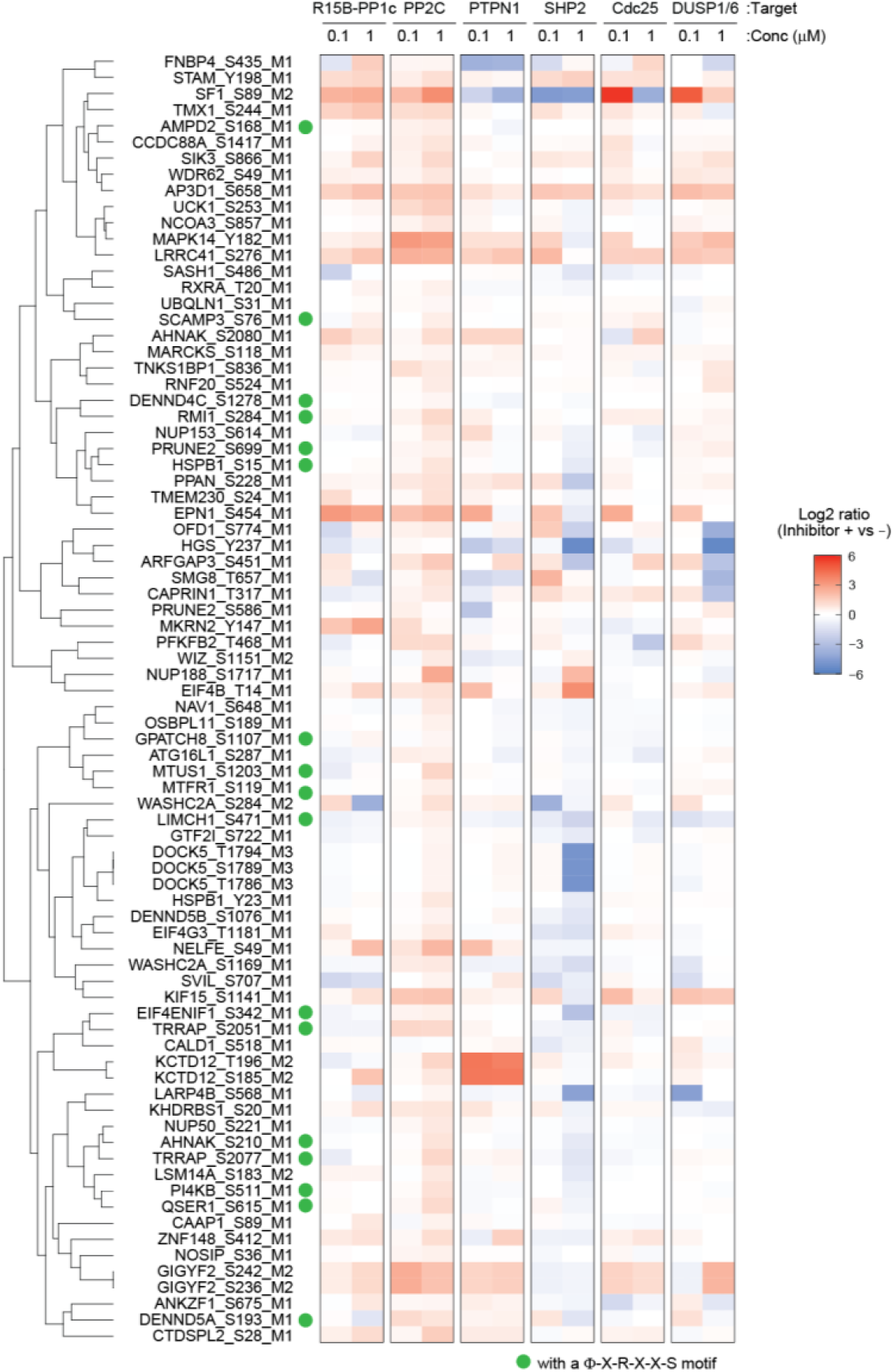
Heatmap showing the log2-transformed phosphosite abundance changes induced by inhibitor treatment.

## References

1. Kholodenko, B. N. Cell-signalling dynamics in time and space. Nat. Rev. Mol. Cell Biol. 7, 165–176 (2006)

2. Lemmon, M. A. & Schlessinger, J. Cell signaling by receptor tyrosine kinases. Cell 141, 1117–1134 (2010)

3. Ardito, F., Giuliani, M., Perrone, D., Troiano, G. & Lo Muzio, L. The crucial role of protein phosphorylation in cell signaling and its use as targeted therapy (Review). Int. J. Mol. Med. 40, 271–280 (2017)

4. Mertins, P. et al. Investigation of protein-tyrosine phosphatase 1B function by quantitative proteomics. Mol. Cell. Proteomics 7, 1763–1777 (2008)

5. Zhang, X. et al. Quantitative phosphoproteomics reveals novel phosphorylation events in insulin signaling regulated by protein phosphatase 1 regulatory subunit 12A. J. Proteomics 109, 63–75 (2014)

6. Rusin, S. F., Schlosser, K. A., Adamo, M. E. & Kettenbach, A. N. Quantitative phosphoproteomics reveals new roles for the protein phosphatase PP6 in mitotic cells. Sci. Signal. 8, rs12 (2015)

7. Kauko, O. et al. Label-free quantitative phosphoproteomics with novel pairwise abundance normalization reveals synergistic RAS and CIP2A signaling. Sci. Rep. 5, 1–17 (2015)

8. Kruse, T. et al. Mechanisms of site-specific dephosphorylation and kinase opposition imposed by PP2A regulatory subunits. EMBO J. 39, e103695 (2020)

9. Batth, T. S. et al. Large-Scale Phosphoproteomics Reveals Shp-2 Phosphatase-Dependent Regulators of Pdgf Receptor Signaling. Cell Rep. 22, 2784–2796 (2018)

10. Vemulapalli, V. et al. Time-resolved phosphoproteomics reveals scaffolding and catalysis-responsive patterns of SHP2-dependent signaling. Elife 10, (2021)

11. Aburai, N., Yoshida, M., Ohnishi, M. & Kimura, K.-I. Sanguinarine as a potent and specific inhibitor of protein phosphatase 2C in vitro and induces apoptosis via phosphorylation of p38 in HL60 cells. Biosci. Biotechnol. Biochem. 74, 548–552 (2010)

12. Krzyzosiak, A. et al. Target-Based Discovery of an Inhibitor of the Regulatory Phosphatase PPP1R15B. Cell 174, 1216–1228.e19 (2018)

13. Ito, Y. et al. Therapeutic effects of the allosteric protein tyrosine phosphatase 1B inhibitor KY-226 on experimental diabetes and obesity via enhancements in insulin and leptin signaling in mice. J. Pharmacol. Sci. 137, 38–46 (2018)

14. Chen, Y.-N. P. et al. Allosteric inhibition of SHP2 phosphatase inhibits cancers driven by receptor tyrosine kinases. Nature 535, 148–152 (2016)

15. Lazo, J. S. et al. Identification of a potent and selective pharmacophore for Cdc25 dual specificity phosphatase inhibitors. Mol. Pharmacol. 61, 720–728 (2002)

16. Korotchenko, V. N. et al. In vivo structure-activity relationship studies support allosteric targeting of a dual specificity phosphatase. Chembiochem 15, 1436–1445 (2014)

17. Easton, J. B., Royer, A. R. & Middlemas, D. S. The protein tyrosine phosphatase, Shp2, is required for the complete activation of the RAS/MAPK pathway by brain-derived neurotrophic factor. J. Neurochem. 97, 834–845 (2006)

18. Ahmad, M. K., Abdollah, N. A., Shafie, N. H., Yusof, N. M. & Razak, S. R. A. Dual-specificity phosphatase 6 (DUSP6): a review of its molecular characteristics and clinical relevance in cancer. Cancer Biol Med 15, 14–28 (2018)

19. Li, C., Scott, D. A., Hatch, E., Tian, X. & Mansour, S. L. Dusp6 (Mkp3) is a negative feedback regulator of FGF-stimulated ERK signaling during mouse development. Development 134, 167–176 (2007)

20. Moon, J., Ha, J. & Park, S.-H. Identification of PTPN1 as a novel negative regulator of the JNK MAPK pathway using a synthetic screening for pathway-specific phosphatases. Sci. Rep. 7, 12974 (2017)

21. Wang, Z., Wang, M., Lazo, J. S. & Carr, B. I. Identification of epidermal growth factor receptor as a target of Cdc25A protein phosphatase. J. Biol. Chem. 277, 19470–19475 (2002)

22. Hanada, M. et al. Regulation of the TAK1 signaling pathway by protein phosphatase 2C. J. Biol. Chem. 276, 5753–5759 (2001)

23. Li, R. et al. Metal-dependent protein phosphatase 1A functions as an extracellular signal-regulated kinase phosphatase. FEBS J. 280, 2700–2711 (2013)

24. Takekawa, M., Maeda, T. & Saito, H. Protein phosphatase 2Cα inhibits the human stress-responsive p38 and JNK MAPK pathways. EMBO J. 17, 4744–4752 (1998)

25. Schaaf, K. et al. Mycobacterium tuberculosis exploits the PPM1A signaling pathway to block host macrophage apoptosis. Sci. Rep. 7, 42101 (2017)

26. Pereira, J. M. et al. Infection Reveals a Modification of SIRT2 Critical for Chromatin Association. Cell Rep. 23, 1124–1137 (2018)

27. Yadav, Y. & Dey, C. S. PP2Cα positively regulates neuronal insulin signalling and aggravates neuronal insulin resistance. FEBS J. 289, 7561–7581 (2022)

28. Batth, T. S. et al. Protein Aggregation Capture on Microparticles Enables Multipurpose Proteomics Sample Preparation. Mol. Cell. Proteomics 18, 1027–1035 (2019)

29. Bekker-Jensen, D. B. et al. A Compact Quadrupole-Orbitrap Mass Spectrometer with FAIMS Interface Improves Proteome Coverage in Short LC Gradients. Mol. Cell. Proteomics 19, 716–729 (2020)

30. Olsen, J. V. et al. Global, in vivo, and site-specific phosphorylation dynamics in signaling networks. Cell 127, 635–648 (2006)

31. Colaert, N., Helsens, K., Martens, L., Vandekerckhove, J. & Gevaert, K. Improved visualization of protein consensus sequences by iceLogo. Nat. Methods 6, 786–787 (2009)

32. Sugiyama, N., Imamura, H. & Ishihama, Y. Large-scale Discovery of Substrates of the Human Kinome. Sci. Rep. 9, 10503 (2019)

33. Breuer, K. et al. InnateDB: systems biology of innate immunity and beyond--recent updates and continuing curation. Nucleic Acids Res. 41, D1228–33 (2013)

34. Szklarczyk, D. et al. The STRING database in 2023: protein-protein association networks and functional enrichment analyses for any sequenced genome of interest. Nucleic Acids Res. 51, D638–D646 (2023)

35. Stokoe, D., Caudwell, B., Cohen, P. T. & Cohen, P. The substrate specificity and structure of mitogen-activated protein (MAP) kinase-activated protein kinase-2. Biochem. J 296 (Pt 3), 843–849 (1993)

36. Manke, I. A. et al. MAPKAP kinase-2 is a cell cycle checkpoint kinase that regulates the G2/M transition and S phase progression in response to UV irradiation. Mol. Cell 17, 37–48 (2005)

37. Pan, C., Olsen, J. V., Daub, H. & Mann, M. Global effects of kinase inhibitors on signaling networks revealed by quantitative phosphoproteomics. Mol. Cell. Proteomics 8, 2796–2808 (2009)

38. Hein, J. B. et al. Phosphatase specificity principles uncovered by MRBLE:Dephos and global substrate identification. Mol. Syst. Biol. 19, e11782 (2023)

39. Choi, J., He, N., Sung, M.-K., Yang, Y. & Yoon, S. Sanguinarine is an allosteric activator of AMP-activated protein kinase. Biochem. Biophys. Res. Commun. 413, 259–263 (2011)

40. Seifen, E., Adams, R. J. & Riemer, R. K. Sanguinarine: a positive inotropic alkaloid which inhibits cardiac Na+,K+-ATPase. Eur. J. Pharmacol. 60, 373–377 (1979)

41. Olsen, J. V. et al. Quantitative phosphoproteomics reveals widespread full phosphorylation site occupancy during mitosis. Sci. Signal. 3, ra3 (2010)

42. Braun, M. et al. DUSP1/6 Inhibition Reduces Tumor Cells and Activates Immune Response in Chronic Lymphocytic Leukemia. Blood 132, 2857 (2018)

